# On the use of generative models for evolutionary inference of malaria vectors from genomic data

**DOI:** 10.1101/2025.06.26.661760

**Authors:** Amelia Adibe Eneli, Pui Chung Siu, Manolo F. Perez, Austin Burt, Matteo Fumagalli, Sara Mathieson

**Affiliations:** School of Biological and Behavioural Sciences, Queen Mary University of London, Mile End Road, E1 4NS, London, United Kingdom; Real Jardín Botánico, CSIC, 2 Pl. Murillo, 28014, Madrid, Spain; Department of Life Sciences, Imperial College London, Silwood Park, SL5 7PY, Ascot, United Kingdom; The Alan Turing Institute, 96 Euston Road, NW1 2DB, London, United Kingdom; Department of Biology, University of Pennsylvania, Philadelphia, Pennsylvania, 19104, USA; Department of Computer Science, Haverford College, Haverford, Pennsylvania, 19041, USA

**Keywords:** malaria parasite, demographic inference, generative adversarial networks, population genetics

## Abstract

Malaria in sub-Saharan Africa is transmitted by mosquitoes, in particular the *Anopheles gambiae* complex. Efforts to control the spread of malaria have often focused on these vectors, but less is known about the demographic history of populations and species in the *Anopheles* complex. Here, we first quantify the genetic structure of mosquito populations in sub-Saharan Africa using unsupervised machine learning. We then adapt and apply an innovative generative deep learning algorithm to infer the joint evolutionary history of populations sampled in Guinea and Burkina Faso, West Africa. We further develop a novel model selection approach and discover that an evolutionary model with migration fits this pair of populations better than a model without post-split migration. For the migration model, we find that our method accurately captures population genetic differentiation. These findings demonstrate that machine learning and generative models are a valuable direction for future understanding of the evolution of malaria vectors, including the joint inference of demography and natural selection. Understanding changes in population size, migration patterns, and adaptation in hosts, vectors, and pathogens will assist malaria control interventions, with the ultimate goal of predicting nuanced outcomes from insecticide resistance to population collapse.

## Introduction

### Genetic monitoring of malaria vectors: challenges and opportunities

Malaria is a tropical disease that, in humans, is mainly caused by the *Plasmodium falciparum* parasite transmitted by the female *Anopheles* mosquito. Infected female *Anopheles* mosquitoes introduce parasites through their saliva into the bloodstream of humans they bite (Milner 2018). The World Malaria Report 2024, released by the World Health Organization (WHO), indicates a rise in malaria cases worldwide, with an estimated 263 million cases in 2023, compared to 252 million in 2022 and 247 million in 2021 (Organization 2024; Poespoprodjo *et al*. 2023). Malaria interventions and elimination efforts have frequently focused on vector control methods. These have been achieved through preventive measures and, where unsuccessful, curative therapies have been used to treat the disease (Phillips *et al*. 2017). More recently, gene drive vector control strategies that employ genetic engineering are being developed to modify mosquito genomes (Nolan 2021). Field release has not yet been performed, but if successful, it would suppress the number of female mosquitoes to promote population collapse.

Genomic surveillance of malaria vectors has gained importance with the emergence and spread of insecticide resistance within and between mosquito populations. Insecticide resistance in mosquitoes is a significant challenge for vector control strategies aimed at reducing mosquito-borne diseases in sub-Saharan Africa. In addition to genetic mechanisms, environmental and operational factors contribute to the development and spread of insecticide resistance in mosquito populations. Environmental factors include local climatic, ecological, agricultural, and urban conditions (Nkya *et al*. 2013; Owusu *et al*. 2017), while operational factors refer to the repeated and improper use of insecticides (Hobbs *et al*. 2023). Genetic resistance to insecticides encompasses target-site, metabolic, cuticular, and behavioural mechanisms. Target-site resistance are changes in proteins targeted by insecticides. A key example is the knockdown mutation (kdr) in the voltage-gated sodium channel gene that confers resistance to pyrethroids and DDT. The kdr mutation occurs in two forms in African Anopheline mosquitoes: kdr-east (L1014S/L995S) and kdr-west (L1014F/L995F) (Grigoraki *et al*. 2021; Mwagira-Maina *et al*. 2021; Suh *et al*. 2023). Metabolic resistance involves changes in genes coding for detoxification enzymes, primarily cytochrome P450s, esterases, and glutathione S-transferases (Adedeji *et al*. 2020). These enzymes can break down insecticides before they reach their target sites. Cuticular resistance refers to genetic changes that reduce insecticide penetration, often associated with thickening of the cuticle or altered compositions (Wood *et al*. 2010). Finally, behavioural resistance enables mosquitoes to avoid contact with insecticides, although this mechanism is still poorly understood (Gatton *et al*. 2013; Carrasco *et al*. 2019).

The extent to which resistance mutations can spread throughout the African continent is still under investigation (Hancock *et al*. 2022). Therefore, inferring adaptive and neutral demographic histories of mosquito populations using genomic data presents an opportunity to improve our understanding of malaria spread. These inferences range from the estimation of the size of past populations and migration rates to the identification of genetic variants under natural selection (Cheng and Steinrücken 2024). A comprehensive understanding of the genetic history of *Anopheles* mosquitoes would inform robust surveillance and control measures against these vectors, the pathogens they spread, and would help predict their potential future demographic changes.

Effective resistance management requires integrated approaches, including rotating insecticides with different modes of action, using synergists to improve insecticide efficacy, and exploring and employing alternative control methods such as biological interventions, e.g., fungi and bacteria (Kamareddine 2012). Continuous monitoring and adaptive strategies are also essential to mitigate the impact of resistance on mosquito control efforts. In order to plan an effective control strategy, it is important to assess the relationship between genetic diversity in *Anopheles* mosquitoes and their dispersal rate and adaptive potential to environmental changes. This includes the estimation of demographic parameters such as the effective population size, population growth rates, and migration between populations. Genetic variation is also shaped by other geographical factors, including levels of urbanisation and trade flows between cities. Understanding the interactions between demographic history, population structure, gene flow, and local selection pressures is critical to predict and manage the spread of insecticide resistance in mosquito populations.

The large-scale genomic resources generated by the MalariaGEN consortium have provided valuable information on the demographic history of *Anopheles*. Using *∂a∂i* (Gutenkunst *et al*. 2009), a coalescent-based approach based on the site frequency spectrum, scientists inferred an expansion in populations located north of the Congo Basin that occurred between 7,000 and 25,000 years ago (Anopheles gambiae 1000 Genomes Consortium 2017). Such expansions could be related to the expansion of human populations as a result of agricultural advances (Li *et al*. 2014). A more recent signal of population bottlenecks, associated with the widespread use of insecticides, was also observed in some populations (Anopheles gambiae 1000 Genomes Consortium 2017).

Other demographic inference methods, such as `SMC++` (Terhorst *et al*. 2017), have been applied to mosquito populations, specifically to the invasive species *Aedes aegypti* (Kent *et al*. 2025). These results also indicate occurrences of population bottlenecks (of varying severity) and re-expansions in the recent past. In addition, dispersal rates and locations of malaria vectors have been inferred using neural networks (Smith *et al*. 2023; Smith and Kern 2023; Battey *et al*. 2020). Due to the complexity of the *Anopheles* system and the ability of neural network-based methods to uncover subtle signals from noisy data, machine learning methods provide a way forward to understand their historical processes.

### Generative models for population genetic data

Supervised machine learning, and deep learning algorithms specifically, have recently emerged as a powerful tool to address some of the most complex questions in population genetics (Schrider and Kern 2018; Korfmann *et al*. 2023). Convolutional neural networks, in particular, have been used for inference of natural selection, changes in population size, variation of recombination rate, and time to the most recent common ancestor, among other processes (Flagel *et al*. 2018; Torada *et al*. 2019). Deep learning has also been shown to outperform traditional simulation-based methods such as Approximate Bayesian Computation (ABC) (Sanchez *et al*. 2021). In the context of studies on malaria, machine learning has been deployed to detect genomic signatures of selection in vectors (Xue *et al*. 2021) and *Plasmodium* parasites (Deelder *et al*. 2021).

Generative models have recently gained popularity, with the ability to create novel text, image, audio, and video from available data. For a given real data set, a generative model refers to any way of quantifying its *distribution*, which can then be used to create synthetic examples in the style of the data set. For biological applications, generative models are an even more recently introduced technology, but they have already shown promising results. In population genetics specifically, simulated data are used extensively for intuition, validation, comparison of methods, and more recently for training machine learning models. Therefore, developing methods to create more realistic simulated data is very desirable. Early generative models in population genetics have been based on generative adversarial networks (GANs) (Wang *et al*. 2021; Yelmen *et al*. 2021, 2023). Yelmen and Jay provided a comprehensive review of generative models for population genetics (Yelmen and Jay 2023).

GANs work by training two models in concert: a generator that creates synthetic examples and a discriminator that predicts whether examples are real or generated. Throughout the training process, both models ideally improve so that, in the end, the generator is creating synthetic examples that confuse the discriminator. In some implementations (Yelmen *et al*. 2021, 2023; Szatkownik *et al*. 2024), the generator is a neural network, creating artificial genetic data with the same Single Nucleotide Polymorphism (SNP) patterns as real data. This is useful for genomic privacy and downstream association studies or polygenic trait analysis. In another study (Wang *et al*. 2021), the generator is based on an evolutionary model, where, during training, the goal is to estimate the parameters of this evolutionary model that create data that closely match the real data (from the perspective of the discriminator). One disadvantage of this method, called `pg-gan`, is that it produces a point estimate of demographic parameters. To better capture uncertainty, Gower and coworkers (Gower *et al*. 2023) developed a variation that uses kernel density estimation to estimate a distribution for each evolutionary parameter, by weighting parameter estimates by their discriminator score (i.e., how realistic are the resulting simulations).

As the discriminator produces a probability (with closer to 1 meaning more “real” and closer to 0 meaning more “fake”), this trained neural network can be used to identify regions of real data containing features unmodeled in the simulations. For example, regions of real data with a very high discriminator score (i.e. very unlike neutral simulations) may be under natural selection or display unusual recombination or mutational features. In a recent study (Riley *et al*. 2024), the `pg-gan` discriminator is fine-tuned using a transfer learning approach to detect various forms of natural selection.

Our motivation for using `pg-gan` is to understand and quantify the demographic history of *Anopheles gambiae* mosquitoes. Specifically, we use `pg-gan` to detect population splits, effective population size changes, exponential growth, and migration rates. Other authors have used `pg-gan` for the inference of demography in mosquito populations (Small *et al*. 2023). As the method was originally developed for and applied to human genetic data, our first objective is to adapt `pg-gan` to take into account the nuances of mosquito data. We then propose and apply a novel method to compare competing historical scenarios and estimate demographic parameters using GANs. GANs are used to fit a series of demographic models, then a separate neural network model is designed to discriminate between them. This network can be used to identify the most probable demographic history for real mosquito data. Finally, we define future research directions on the development and use of generative models in population genetics for pathogens and disease-vectors. Our software, called `pg-gan-mosquito`, is open-source and available at https://github.com/mathiesonlab/pg-gan-mosquito/.

## Materials and Methods

### Genetic data and model exploration

The *Anopheles gambiae* complex is composed of at least eight mosquito species, five of which are the main malaria vectors: *An. gambiae, An. coluzzii, An. arabiensis, An. melus* and *An. merus* (Charlwood 2019). To assess their genetic diversity, population structure and demographic history, we analysed genomic data of samples captured in 13 sub-Saharan African countries of two species of the complex, i.e., *Anopheles gambiae, Anopheles coluzzii*, and three types of hybrid strains from the Ag1000G Phase 2 open access dataset (Anopheles gambiae 1000 Genomes Consortium 2024), for a total sample size of *n* = 1142 mosquitoes. Following suggested data filtering (Anopheles gambiae 1000 Genomes Consortium 2017), we assessed population structure using Uniform Manifold Approximation and Projection (UMAP) (Diaz-Papkovich *et al*. 2021).

To show the applicability of `pg-gan-mosquito` to this system, we focused on a pair of populations, GN (Guinea) and BF (Burkina Faso), due to their genetic similarity and geographic proximity. We retrieved haplotype data from 112 *An. gambiae* samples (31 from Guinea (GN) and 81 from Burkina Faso (BF)) from the MalariaGEN database, Phase 2, following a similar data processing protocol to a previous study (Anopheles gambiae 1000 Genomes Consortium 2017). Specifically, we considered only biallelic variants in chromosomal arms 3L and 3R that passed quality control and were located in heterochromatin states. Furthermore, we filtered out SNPs within the *Gste* gene region, a known target of selection associated with insecticide resistance.

For our simulations, we assumed a mutation rate of 3.5 *×* 10^*−*9^ per site per generation, with 11 generations per year (Anopheles gambiae 1000 Genomes Consortium 2017), and a recombination rate of 1.45 *×* 10^*−*8^ per site per generation (Adrion *et al*. 2020; Lauterbur *et al*. 2023). We sought to compare our demographic inferences with previously estimated models (Anopheles gambiae 1000 Genomes Consortium 2017) that incorporated only information from the joint site frequency spectrum using the software *∂a∂i* (Gutenkunst *et al*. 2009). Using the ensemble of such previous models, we computed the median for each parameter, as the model with the lowest Akaike Information Criterion (AIC) score was sometimes an outlier. We refer the median for demographic models with and without bidirectional migration (Anopheles gambiae 1000 Genomes Consortium 2017) as *mig* and *no-mig*, respectively.

### Generative adversarial model

To assess GAN training, we should observe the following patterns in loss functions and discriminator accuracy. At the start of training, the discriminator should correctly identify real data from simulations, resulting in high accuracy for both types of data. Here, accuracy is measured as the fraction of regions correctly identified as real or simulated data. The loss of the discriminator should be low (easily able to distinguish real vs. simulated), and the loss for the generator should be high (since its generated data are not “fooling” the discriminator yet). As training progresses, the generator improves the quality of the simulations and reduces its loss, and the discriminator is progressively having difficulties distinguishing real data from simulated data. During the stable competing phase the loss functions have more or less plateaued, although there may still be minor fluctuations in the parameter choices of the generator. Ideally, at the end of the training, the discriminator accuracy is around 50% for both the real and the simulated data, indicating high quality simulations that confuse the discriminator.

To adapt `pg-gan` for mosquito genetic data, we made several modifications to the hyper-parameters and training procedure. We used the `AdamW` optimizer instead of the regular Adam optimizer for robust convergence. We changed the number of units in the first fully connected layer to 160. During pre-training (an initial phase of parameter exploration to make sure we are starting from a successful discriminator), we set the dropout rate to 0.5 and the learning rate to 1 *×* 10^*−*3^, and during the main training loop we used a dropout rate of 0.8 and a learning rate of 25 *×* 10^*−*6^. Since mosquitoes have a higher density of genetic variants than humans, we increased the number of SNPs per region from 36 to 72 (to still preserve computational feasibility). Since we have a different sample size in each population, we change the permutation-invariant collapsing function from `sum to mean`.

If a successful training run is not observed (see Figure S1 for an example) we created several debugging strategies. During pre-training, the goal is to try random parameter combinations to find an initial configuration that produces simulated data that is easily distinguishable from real data. If the number of pre-training iterations is reached without reaching a sufficient discriminator accuracy, we can a) increase the parameter search space, b) increase the model capacity by adding more filters or hidden units, and/or c) lower the dropout rate (while being aware of over-fitting). If the discriminator quickly obtains a high accuracy, the dropout rate can be increased, or we force the pre-training to continue.

During training, if the discriminator is over-fitting (indicated by very high accuracy) we can reduce the number of training iterations. `AdamW` also reduces the magnitude of the weights which helps with over-fitting. If the discriminator is under-fitting, we increase the model capacity and/or lower dropout rate. Finally, if the discriminator accuracy remains high after pre-training, this could suggest that the evolutionary model is not expressive enough to create simulations matching the real data. A different or more complex evolutionary model is likely needed in these circumstances.

Using this modified version of `pg-gan`, we fit two different demographic models. The first model (Anopheles gambiae 1000 Genomes Consortium 2017) (*no-mig*) begins with an ancestral population of size *N*_*I*_ . At time *T*_*G*_ the ancestral population can start to grow or contract (exponentially), with a final size of *N*_*F*_ right before splitting. The population split occurs at *T*_*S*_, with population sizes *N*_*I*1_ and *N*_*I*2_ for the two populations, respectively. Both populations can undergo exponential size changes until their final sizes, *N*_*F*1_ and *N*_*F*2_. The second model (*mig*) has the same structure, but with bidirectional migration after the split (parameter *M*_*G*_ ).

### Demographic model selection and evaluation

To assess which demographic model is a better fit to the data, we developed a novel machine learning-based model selection algorithm inspired by previous studies (Fonseca *et al*. 2021; Kirschner *et al*. 2022). We trained an additional neural network (with the same CNN architecture as the `pg-gan-mosquito` discriminator) with data simulated under both models. The task of this network is to perform a binary classification (whether or not each region comes from the *no-mig* or *mig* model). After training is complete, we feed regions of *real* data into the trained network to predict whether their histories are more similar to the *no-mig* or *mig* model. At the end of this step we have the fractions of real regions assigned to each model, which allows us to specify which model is better for the real data and how confident we are about this prediction.

For the better fitting model, we also sought to access whether data simulated under this model resembles real data. To do this, we compared the distributions of commonly used summary statistics of genetic diversity and differentiation (Nielsen 2005), as previously done (Wang *et al*. 2021). These summary statistics include pairwise heterozygosity (*π*), Watterson’s *θ*, site frequency spectrum, inter-SNP distance, linkage disequilibrium (LD) decay, number of haplotypes and a measure of population genetic differentiation (*F*_*ST*_ ).

## Results

After using UMAP to assess the genetic structure in the *Anopheles* complex (Anopheles gambiae 1000 Genomes Consortium 2020), we confirm that the samples tend to cluster closely according to their species and country of origin (Figure S2). The mainland populations are tightly clustered in both groups with hybrid forms on the periphery of the *An. coluzzii* cluster. Similarly, *An. gambiae* samples from mainland populations form a large cluster, with the different populations appearing distinctly. Samples from Gabon, Uganda, and the Mayotte island form separate and distant clusters. To understand the joint demographic history and divergence of the GN (Guinea) population and the BF (Burkina Faso) population, we fit two different demographic models using `pg-gan-mosquito` – *no-mig* (no migration after the population split) and *mig* (migration after the population split). Figures 1 and 2 show successful training runs for the *no-mig* and *mig* models, respectively. In both cases, we observe that the generator and discriminator losses are well-matched by the end of training. For the accuracies, initially the discriminator is easily able to distinguish real from simulated data, but during training it exhibits some difficulty. At the end of training, the *no-mig* model discriminator displays high, but not perfect, accuracy, indicating an appropriate level of confusion compatible with a successful training. Figure S1 shows an example of a failed training run, where learning stops, leading to a plateau in loss and accuracy. This type of failed run is not typical (less than 10% of cases) and strategies for avoiding such outcomes are described below.

**Figure 1.**
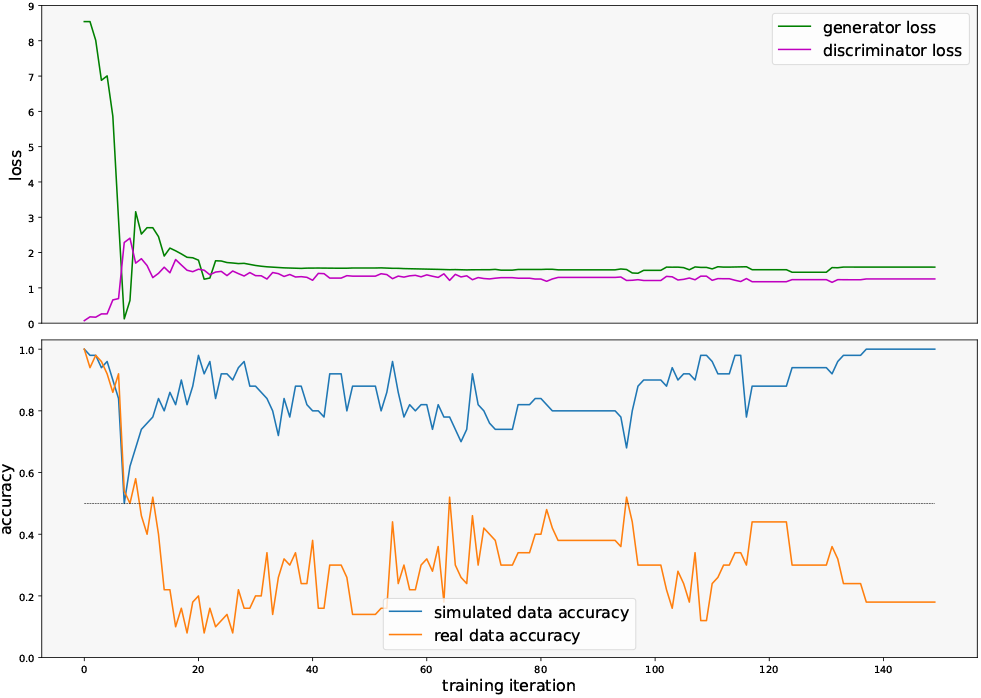
Training run with populations GN and BF under the *no-mig* model. The top panel shows the generator (green) and discriminator (pink) loss functions, which are well-matched by the end of training. The bottom panel shows the discriminator accuracy: blue for simulated data and orange for real data. Accuracy is measured as the proportion of regions the discriminator correctly identifies.

**Figure 2.**
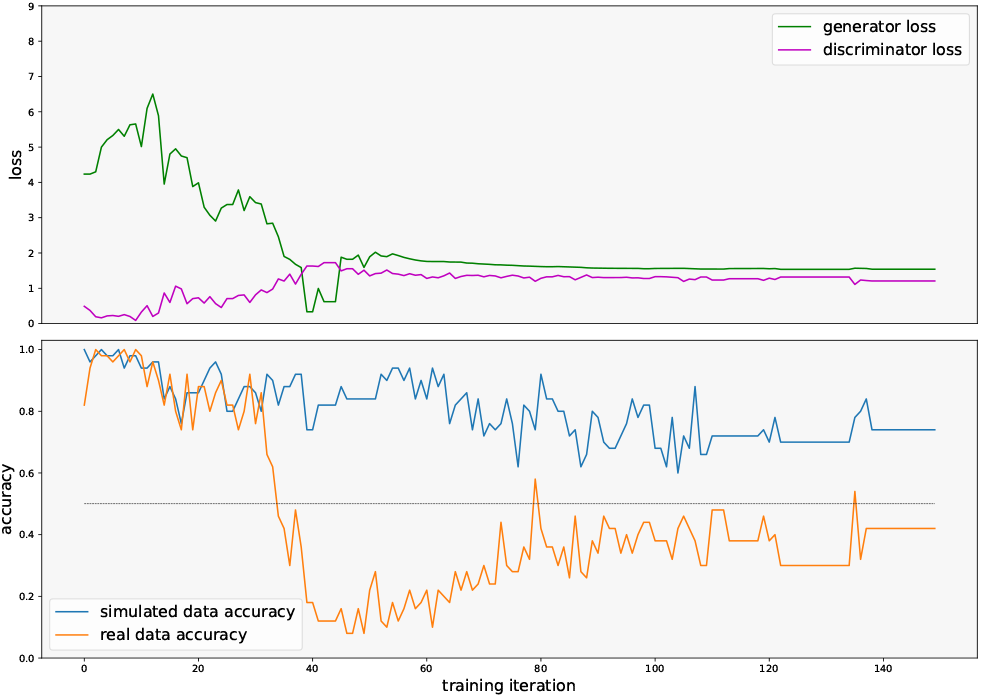
Training run with populations GN and BF under the *mig* model. The top panel shows the generator (green) and discriminator (pink) loss functions, which are well-matched by the end of training. The bottom panel shows the discriminator accuracy: blue for simulated data and orange for real data. Accuracy is measured as the proportion of regions the discriminator correctly identifies.

Table 1 shows the point estimates for the model parameters. For both models, we infer a very recent split time (*T*_*S*_) with large recent population sizes (*N*_*F*1_ and *N*_*F*2_).

**Table 1.** Joint population model parameter inference from `pg-gan-mosquito`. Model parameter descriptions and baseline inference results are based on (Anopheles gambiae 1000 Genomes Consortium 2017). Time units are in generations, *M*_*G*_ = 2*N*_*f*_ *m* where *m* is the bidirectional fractional migration rate per generation.

| name | description | baseline *no-mig* | pg-gan-mosquito *no-mig* | baseline *mig* | pg-gan-mosquito *mig* |
| --- | --- | --- | --- | --- | --- |
| $N_I$ | initial ancestral size | 420,699 | 579,516 | 415,254 | 560,796 |
| $T_G$ | time of size change of ancestral population | 89,497 | 75,249 | 91,234 | 57,254 |
| $N_F$ | size of ancestral population before split | 9,449,179 | 2,784,452 | 9,164,256 | 7,772,061 |
| $T_S$ | time of split | 2,243 | 3,519 | 2,996 | 2,265 |
| $N_{I1}$ | initial size of population GN | 18,328,570 | 117,287,221 | 22,149,704 | 16,137,900 |
| $N_{I2}$ | initial size of population BF | 39,242,967 | 62,771,288 | 11,679,596 | 20,976,274 |
| $N_{F1}$ | final size of population GN | 42,056,997 | 59,092,640 | 31,103,040 | 134,237,900 |
| $N_{F2}$ | final size of population BF | 42,050,613 | 189,066,655 | 19,166,216 | 218,082,996 |
| $M_G$ | migration rate | n/a | n/a | 20 | 27 |

To select a demographic model, we used our model selection approach as described in the Methods. For this approach, we train a new CNN using simulated regions from both models. An accuracy curve is shown in Figure 3, where we plot the training and validation accuracy (both based on simulated data) throughout the training iterations. Both demographic models produce similar data, as demonstrated by the accuracy starting at around 50% and plateauing between 65-70%.

**Figure 3.**
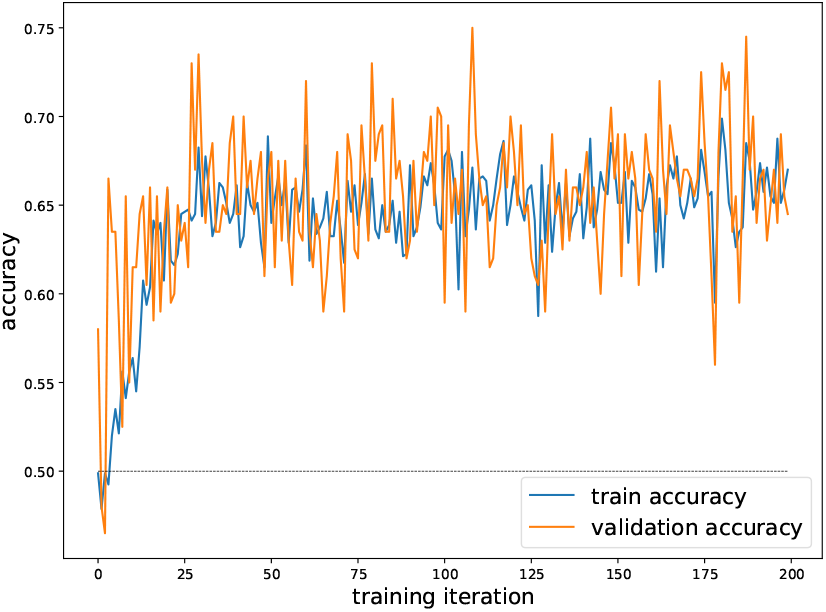
Model selection accuracy curve. We train a discriminator to distinguish data simulated under two models (*no-mig* and *mig*). The accuracy on training and validation data is shown over the training iterations. Although the training is successful, the final accuracy is not very high, indicating these demographic models produce similar data.

**Figure 4.**
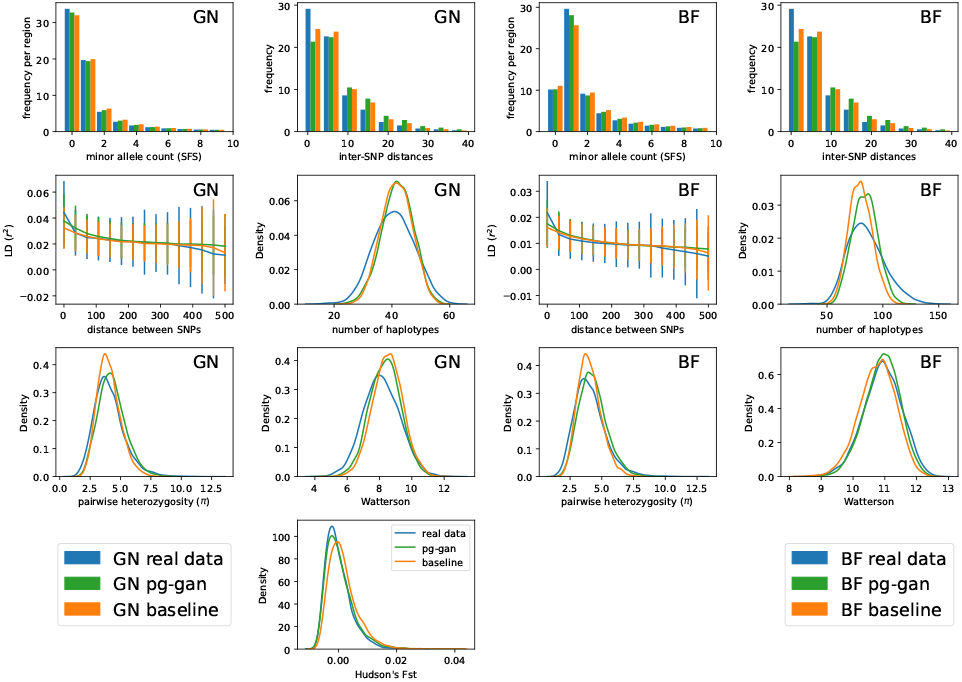
`pg-gan-mosquito` results for GN and BF populations, fitting a demographic history with migration. Summary statistic distributions are shown for three datasets. Blue: real data from either the GN or BF population. Green: simulations under the parameters inferred by `pg-gan-mosquito` inference. Orange: simulations under the parameters inferred by *∂a∂i* (baseline results from (Anopheles gambiae 1000 Genomes Consortium 2017)).

**Figure 5.**
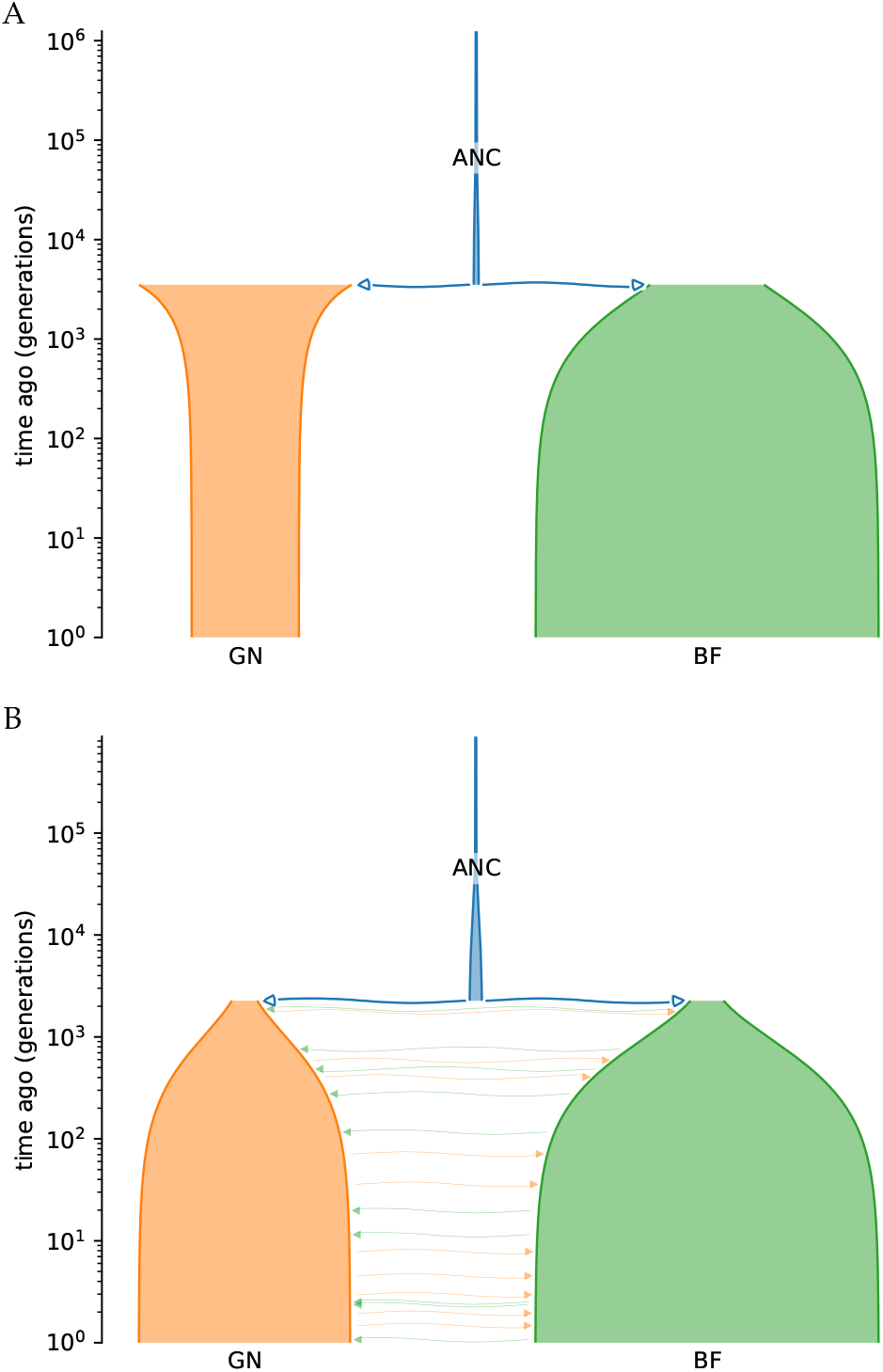
A: `pg-gan-mosquito` inferred demography for GN and BF using a *no-mig* model. B: `pg-gan-mosquito` inferred demography for GN and BF using a model with migration (the more likely history based on our model selection results).

After running the GN/BF real data through the model selection discriminator, the fraction of real regions classified as *mig* model is 0.744, while the remaining 0.256 is classified as *no-mig*. These findings are in line with the observation that these two populations are geographically close. Therefore, we conclude that a model with bidirectional migration explains the observed genetic data better than a model without migration.

To visually assess the match between the data simulated under the inferred parameters of the *mig* model and the real data, we generate distributions of various classical summary statistics. We also plot these distributions under the previously inferred demographic model (hereafter named *baseline*)(Anopheles gambiae 1000 Genomes Consortium 2017). To obtain each distribution, we simulate many regions under each set of parameters (for the simulated data) and sample random regions from the genome (for the real data). Note that SNPs for each region are a combined set of GN and BF haplotypes. When computing summary statistics for the haplotypes from one of these population, we will observe non-segregating sites (i.e. 0 entry in the site frequency spectrum, SFS). These sites indicate the variation was in the other population (or non-segregating in both with different alleles). Figures S3 and 4 show these distributions for the *no-mig* and *mig* model, respectively. For the *mig* model, we generally find a similar match between our results (green) and the real data (blue) to the match between the *baseline* model (orange) and the real data (blue).

To quantify how well summary statistics from simulated data match the real data, we compute Wasserstein distances between summary statistic distributions. These values are provided for both *baseline* model and our estimates using `pg-gan-mosquito` in Table S1 for the *no-mig* model, and Table S2 for the *mig* model. For the *no-mig* model we observe that the data simulated under the *baseline* model are generally closer to the real data. On the other hand, for the *mig* model, data simulated under the `pg-gan-mosquito` model are on average as close to the real data as data simulated under the *baseline* model. We note that *F*_*ST*_ is poorly fitted by the *baseline* model, possibly due to the migration rate being capped to its upper limit 20 (Anopheles gambiae 1000 Genomes Consortium 2017). We acknowledge that migration rates are notoriously difficult to estimate (Gower *et al*. 2023), and we also anticipate some uncertainty in our estimates.

We note that for this migration model, our estimate of *T*_*S*_ (time of population split) is much lower than the compared baseline model, and we retrieve a higher migration rate (although the baseline model was capped at 20, likely hindering their inference). Given the close proximity of these two populations, we believe that a more recent split time with more migration is compatible with our prior expectations based on the biology and ecology of the species. Additionally, the larger recent effective population sizes could account for the observed diversity between (and within) the two populations.

## Discussion

In this study, we refine a generative adversarial network algorithm (`pg-gan` (Wang *et al*. 2021)) for use with genetic data from malaria vectors. We apply this demographic inference method to a pair of mosquito populations (GN from Guinea and BF from Burkina Faso) and fit two different evolutionary models, one with post-split migration and one without. We then develop a model selection method based on discriminating between datasets generated under each model, with the goal of identifying which evolutionary history best fits the real data. Our results indicate a model that includes post-split migration between the GN and BF is most appropriate, consistent with the close geographic proximity of these two populations. Furthermore, the parameters of our fitted demographic history produce simulated data that are as close to the real data as a previously reported result (Anopheles gambiae 1000 Genomes Consortium 2017).

One limitation of the current implementation of `pg-gan-mosquito` is that the input must take the form of biallelic SNPs. Although most variants in the *Anopheles* complex are biallelic, we find that on average approximately 23% of sites are triallelic and approximately 3% are tetrallelic. In particular, multiallelism (i.e. the presence of more than two alleles at a single genetic locus) appears to be highly prevalent in populations that have been sampled near water resources (Figure S4). Swampy vegetation, such as mangroves, may support the lifecycle development (i.e. eggs, larvae, pupae stages) of the mosquitoes as the water entities of such habitats are less disturbed by anthropogenic activities due to the pneumatophores of the surrounding mangrove vegetation. This, most likely in combination with other pollutants (e.g. agrochemicals) in the waters, could potentially promote or accelerate the development of insecticide resistant phenotypes (Richards *et al*. 2020; Williams and Hill 2019). However, it remains unknown whether insecticide resistance is specifically directly correlated with multiallelism (Clarkson *et al*. 2021; Corbel and N’Guessan 2013). Multiallelism is less pronounced in East and West African populations (relative to Central African populations), where bottlenecks or recent colonisation events may have reduced rare variation. In general, future demographic analyses should include information on multiallelic genetic variants to further elucidate recent historical events that affect diversity levels.

There are many directions for future work in this area, including scaling machine learning-based demographic inference methods to more pairs of populations or groups of populations. Currently, data filtering is performed to retain neutral variation only, but natural selection is still likely to impact our results. Future work could incorporate a distribution of fitness effects, as previous studies aimed to infer demographic histories in the presence of selection (Johri *et al*. 2021, 2023; Marsh and Johri 2024). Sites under natural selection could also be identified using post-hoc analyses of outlier regions that do not fit the inferred demographic history (Riley *et al*. 2024). A recently proposed and interesting direction is to incorporate spatial features to infer selection targets in malaria vectors (Rehmann *et al*. 2025). Domain adaptation and transfer learning could also be used to mitigate source/target data mismatch in cases where models have been trained on different species/populations or datasets (Cobb and Smith 2025; Arnab *et al*. 2025). These techniques could also reduce energy consumption by reducing simulation and training time.

Here we used UMAP to understand the population structure of malaria vectors, but complementary approaches such as contrastive learning (Thor and Nettelblad 2025), hierarchical soft clustering (Burger *et al*. 2024), or other supervised non-linear approaches (Qin *et al*. 2022) could augment our analyses. Approaches that compare different machine learning methods could help refine the set of evolutionary histories that best explain the data, leading to more robust conclusions about ongoing and future contact between populations.

We envisage that our new implementation, `pg-gan-mosquito`, will be applicable to a wider range of species with high genetic diversity. By successfully modifying the original implementation, we demonstrate how, in general, generative models are a valuable approach for the demographic inferences of non-model species from genomic data. Recent studies have shown that simulation-based approaches can be a suitable framework for inference of evolutionary histories of malaria parasites (Lefebvre *et al*. 2025), and we argue that our study lays the foundation for future research directions on the joint inference of host, vector and pathogen coevolution using deep learning and synthetic data. Looking ahead to intervention strategies, realistic models of mosquito dispersal and migration between populations could help anticipate the trajectory and tempo of insecticide resistance. A fitted evolutionary history (such as that produced by `pg-gan-mosquito`) provides a null model for selection scans, which could illuminate which genes contribute to insecticide resistance. Finally, knowledge of the relative effective population sizes in different geographic regions could help us predict the effectiveness of gene drive release strategies and the pace of population collapse.

## Data availability

Mosquito data for the project can be accessed through the MalariaGEN project https://www.malariagen.net/, Phase 2 data. Our software, `pg-gan-mosquito` is open-source and available on GitHub: https://github.com/mathiesonlab/pg-gan-mosquito/. We also provide instructions on how to use `pg-gan-mosquito`, which helps researchers of other species who are interested in generative models, demographic inference, and selection of evolutionary models.

