## Supplementary Material for "On the use of generative models for evolutionary inference of malaria vectors from genomic data"

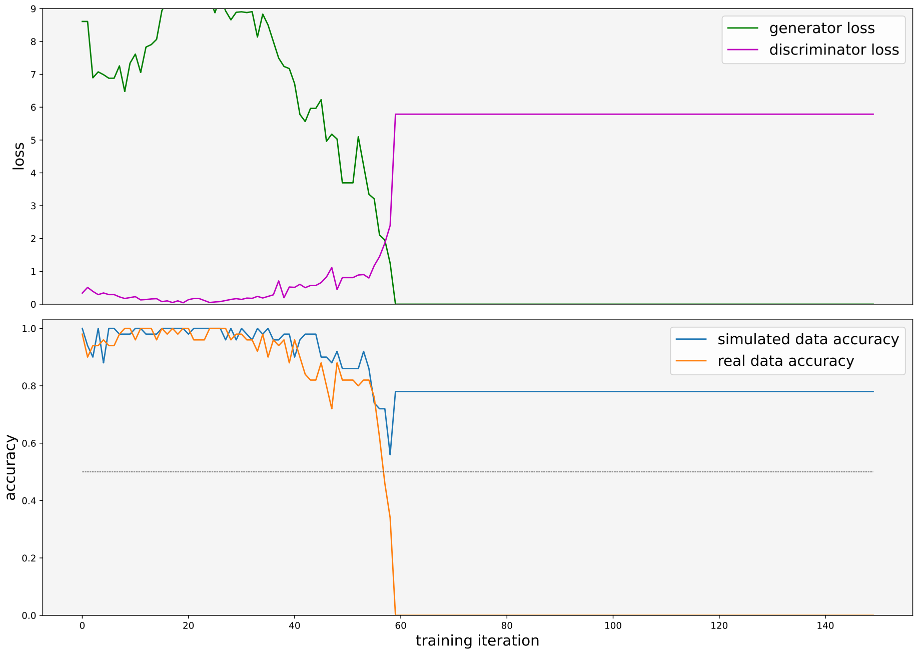


Figure S1: `pg-gan` training run with populations GN and BF under the *mig* model, where training was unsuccessful. The top panel shows the generator (green) and discriminator (pink) loss functions. The bottom panel shows the discriminator accuracy: blue for simulated data and orange for real data. Around training iteration 60, learning stops and the model weights do not change, leading to a plateau in both losses and both accuracies.

| summary statistic | `pg-gan-mosquito` | baseline |
|---|---|---|
| GN minor allele count (SFS) | 0.6529 | **0.3615** |
| GN inter-SNP distances | 7.5684 | **5.0228** |
| GN distance between SNPs | 0.0063 | **0.0017** |
| GN number of haplotypes | 3.683 | **1.4876** |
| GN pairwise heterozygosity ($\pi$) | 1.2472 | **0.2023** |
| GN Watterson | 0.4592 | **0.3015** |
| BF minor allele count (SFS) | 0.6318 | **0.5672** |
| BF inter-SNP distances | 7.5684 | **5.0228** |
| BF distance between SNPs | 0.0028 | **0.0009** |
| BF number of haplotypes | 7.5718 | **5.0602** |
| BF pairwise heterozygosity ($\pi$) | 1.2429 | **0.2082** |
| BF Watterson | 0.0501 | **0.041** |
| GN/BF Hudson's $F_{ST}$ | **0.0004** | 0.0007 |

Table S1: Wasserstein distances for the *no-mig* model. The left-hand column shows common population genetic summary statistics, computed for each mosquito population separately (GN and BF) and for $F_{ST}$ together. The middle column is the Wasserstein distance between the summary statistic distributions for the real data and data simulated under the `pg-gan-mosquito`-inferred demography. The last column is the Wasserstein distance between the summary statistic distributions for the real data and data simulated under the *∂a∂i baseline* model. The smaller distance in each row is bolded, indicating the closer match to the real data.

| summary statistic | `pg-gan-mosquito` | baseline |
|---|---|---|
| GN minor allele count (SFS) | **0.2343** | 0.4134 |
| GN inter-SNP distances | 4.96 | **4.1014** |
| GN distance between SNPs | 0.0026 | **0.0019** |
| GN number of haplotypes | 1.6676 | **1.611** |
| GN pairwise heterozygosity ($\pi$) | 0.2646 | **0.2046** |
| GN Watterson | **0.2451** | 0.3864 |
| BF minor allele count (SFS) | **0.3449** | 0.797 |
| BF inter-SNP distances | 4.96 | **4.1014** |
| BF distance between SNPs | 0.0012 | **0.0011** |
| BF number of haplotypes | **4.1946** | 5.9824 |
| BF pairwise heterozygosity ($\pi$) | 0.2568 | **0.2115** |
| BF Watterson | **0.0436** | 0.1575 |
| GN/BF Hudson's $F_{ST}$ | **0.0005** | 0.0016 |

Table S2: Wasserstein distances for the *mig* model. The left-hand column shows common population genetic summary statistics, computed for each mosquito population separately (GN and BF) and for $F_{ST}$ together. The middle column is the Wasserstein distance between the summary statistic distributions for the real data and data simulated under the `pg-gan-mosquito`-inferred demography. The last column is the Wasserstein distance between the summary statistic distributions for the real data and data simulated under the *∂a∂i baseline* model. The smaller distance in each row is bolded, indicating the closer match to the real data.

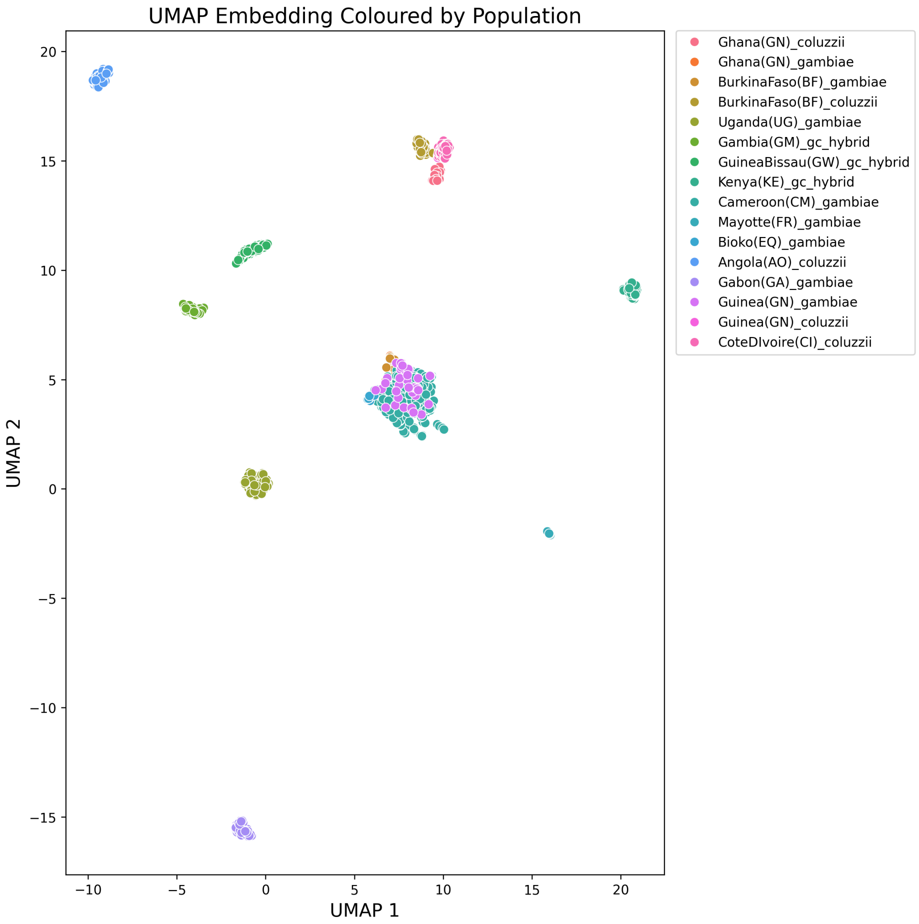


Figure S2: Population structure analysis of mosquitoes using the UMAP technique. Each mosquito represented by a marker. Biallelic SNP data from euchromatic regions of Chromosome arm 3L were projected onto two components denoted as UMAP 1 and UMAP 2.

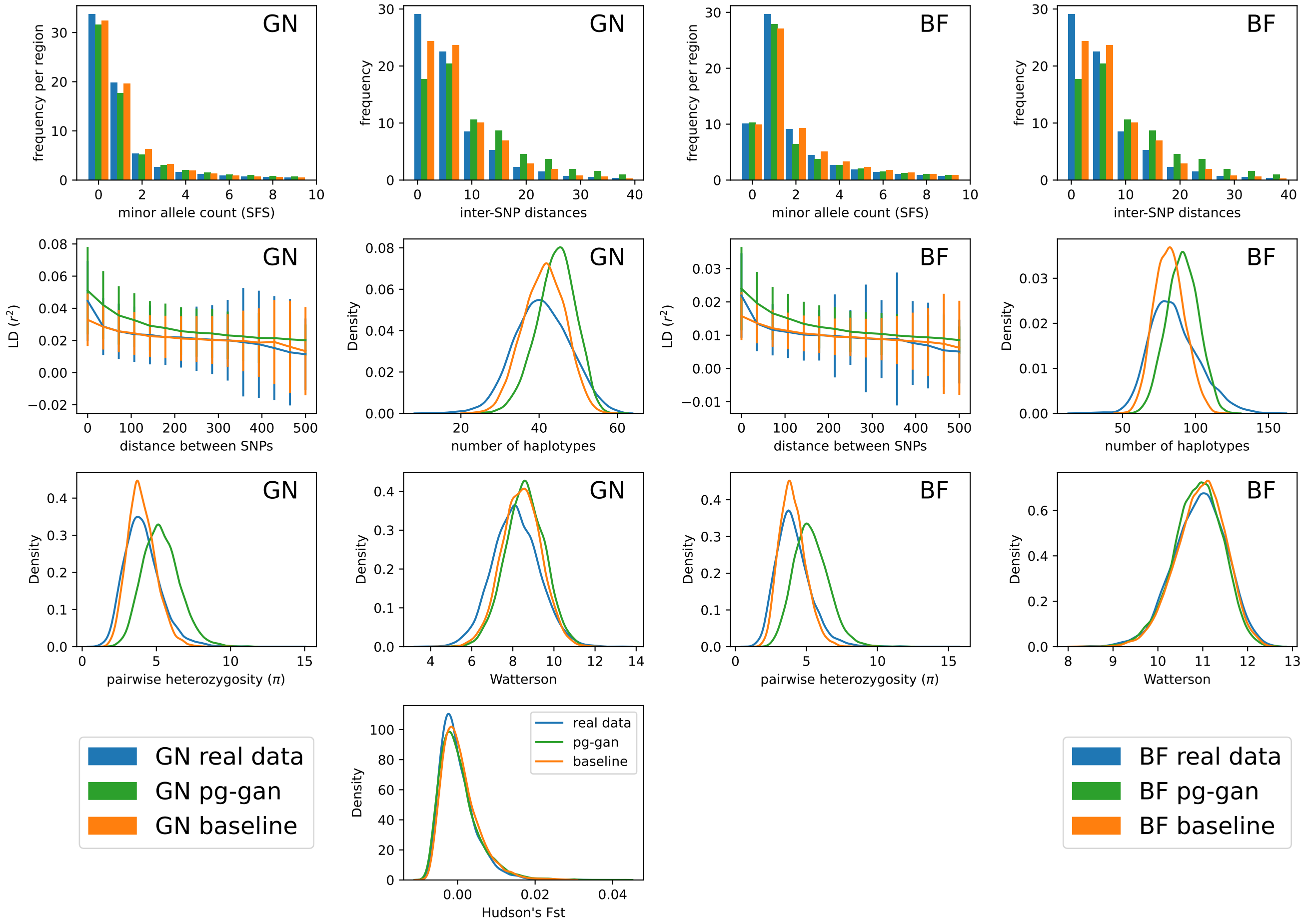


Figure S3: `pg-gan-mosquito` results for GN and BF populations, fitting a demographic history without migration (*no-mig*). Summary statistic distributions are shown for three datasets. Blue: real data from either the GN or BF population. Green: simulations under the parameters inferred by `pg-gan-mosquito` inference. Orange: simulations under the parameters inferred by $\partial a \partial i$ (baseline).

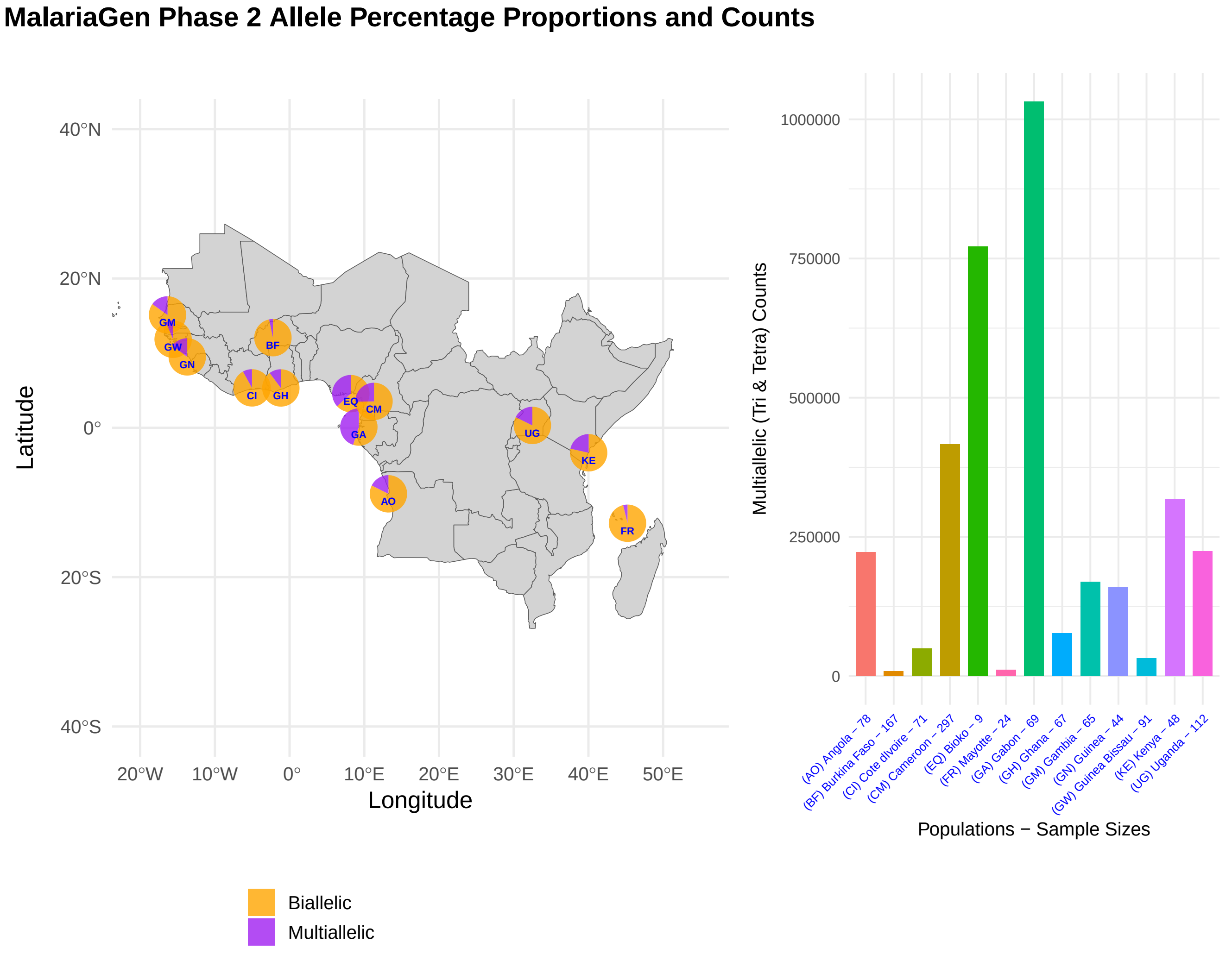


Figure S4: Ag1000G Phase 2 mosquito sample locations. Circle colours denote biallelic (orange) and multiallelic (purple) for triallelic and tetrallelic percentage proportions of each population dataset. The bar chart shows the multiallelic counts for each population.
